# Developing a sensitive indoor air surveillance approach for nosocomial pathogens and antimicrobial resistance

**DOI:** 10.64898/2026.08.28.747720

**Authors:** Sophie Chen, Xenia Kostoulias, Parth Sharma, Chris Greening, Anton Peleg, Rachael Lappan

**Affiliations:** Department of Microbiology, Biomedicine Discovery Institute, Monash University, Clayton, Victoria, 3800, Australia; Department of Infectious Diseases, The Alfred Hospital and School of Translational Medicine, Monash University, Melbourne, Victoria, 3004, Australia; Centre to Impact AMR, Monash University, Clayton, Victoria, 3800, Australia

## Abstract

The role of bioaerosols in the transmission of pathogens and antimicrobial resistance (AMR) is of increasing clinical importance, particularly in settings housing vulnerable populations. Air filtration (e.g. HEPA filtration) and ventilation (e.g. minimum air changes per hour) measures are designed to restrict the airborne transmission of microorganisms. Despite these measures, airborne transmission remains a persistent issue in hospitals, workplaces, aged care, and schools, and is not typically assessed in routine surveillance for infection prevention. Here, we evaluated the efficacy of a high-volume air sampling approach to capture the indoor ‘aerobiome’, and investigated the potential for bioaerosols to mediate disease and AMR transmission in workplace and hospital settings. Our sampling approach demonstrates the benefits of simple decontamination procedures and personal protective equipment on the ability to distinguish genuine low biomass signals in air samples from blank controls, enabling reliable and sensitive microbial detection down to a limit of 69 bacterial cells/m^3^ of air. In a workplace bathroom setting, increased airborne biomass was strongly associated with human activity. This diminished significantly after a few hours of no activity, yet persisted in the indoor environment, with viable identical bacterial strains recovered from bioaerosols and bathroom surfaces across months of sampling. Applying our approach in a hospital ward, air samples from occupied patient rooms were not distinguishable from blank controls and contained negligible fungal and bacterial content, with only trace contributions from human occupancy. Our findings indicate that air filtration measures in this ward are effective at minimising airborne risks, but periodic testing of high-risk areas may be valuable in indoor settings with greater human traffic and may contribute key information to outbreak investigations.

## Introduction

Indoor air is an important transmission route for both pathogens and antimicrobial resistance (AMR), particularly in hospital environments. Aerosolised particles containing microorganisms are generated through human respiratory activities such as coughing and sneezing, during medical procedures including intubation and ventilation, and also via cleaning, toilet flushing, and handwashing.^1–3^ Certain areas may be subject to increased dissemination of respiratory or enteric pathogens when patients present with symptoms such as vomiting, diarrhoea, coughing, or sneezing that can aerosolise infectious bodily fluids. This risk may be further amplified by the commonality of non-lidded toilets and commodes in hospital settings.

The ESKAPE pathogens (*Enterococcus faecium, Staphylococcus aureus, Klebsiella pneumoniae, Acinetobacter baumannii, Pseudomonas aeruginosa, Enterobacter* spp.; sometimes *Escherichia coli*) are the leading causes of healthcare-acquired infections globally and are increasingly multi-drug resistant.^4^ The spread of acquired AMR is a major driver behind the global AMR crisis, particularly via the horizontal transfer of resistance genes mediated by mobile genetic elements. Monitoring both clinically important pathogens and high-risk mobile AMR determinants therefore provides a more complete picture of existing and potential risks. While targeted pathogen detection approaches like quantitative PCR (qPCR) are sensitive and well suited to low-biomass samples such as surface swabs or air samples, they require prior knowledge for target selection. This limits their usefulness in proactive surveillance aimed at preventing unexpected outbreaks in a variety of settings: broader, untargeted monitoring of bioaerosols (the aerobiome) offers the potential to detect new risks and the movement of resistance genes through environments such as hospitals, workplaces, aged care residences, and schools, enabling early interventions.

Previous research on bioaerosol monitoring in hospitals has been effective and informative for infection prevention and control, but has focused heavily on respiratory viruses including SARS-CoV-2. Management of airborne risks relies primarily on filtration and ventilation. These controls include HEPA filtration, UV sterilisation, control of temperature and humidity to minimise droplet lifespan and increase settling,^5,6^ and differential pressure systems designed to prevent airflow out of contaminated environments into cleaner ones. Negative pressure rooms are used to isolate infectious patients, and positive pressure rooms such as operating theatres maintain sterility. Despite these measures, bioaerosols, especially small (1-10 µm) droplets, resist evaporation and contaminate ventilation systems,^7^ and SARS-CoV-2 viral particles have been shown to persist in the hospital environment of infected patients despite extensive ventilation with 12 or more air changes per hour.^8,9^ Surveillance of non-viral pathogens in bioaerosols is less common, though several efforts have been made to characterise AMR and resistomes in hospital air. Antimicrobial resistance gene (ARG) diversity in hospital bioaerosols includes markers for erythromycin, tetracycline, vancomycin, β-lactam, and carbapenem resistance.^10–13^ However, the clinical significance of these aerosolised ARGs remains unclear, as ARG detection often remains disconnected from an understanding of health risk.

Airborne surveillance in hospital environments is not routine. The direct assessment of airborne risks is challenging relative to building an understanding of risk from active patient infections, as air contains much lower biomass than clinical or even other types of environmental samples (such as handrail, sink or toilet swabs). This poses a dual challenge, where air samples may contain too little biomass for accurate surveillance, and are additionally prone to contamination with other microbes during sampling or analysis. This is an ongoing yet increasingly recognised issue within low biomass microbial DNA studies,^14,15^ requiring stringent sampling and handling procedures and careful interpretation of microbial findings alongside equivalent blank control samples. Here, we develop and evaluate an air sampling and detection approach designed to maximise biomass recovery while minimising sample contamination. Using SASS 3100 air samplers previously shown to effectively capture aerosolised SARS-CoV-2 in hospital environments,^9,16^ we combine a high-volume filtration approach, with culture-based validation and microbiome sequencing to assess the feasibility of broadly characterising the indoor aerobiome to detect airborne bacterial and AMR risks before outbreaks occur, providing a generalisable platform for both routine and high-risk surveillance in a variety of settings.

## Results

### Indoor sampling with the SASS 3100 dry air sampler is sensitive and informative

The SASS 3100 dry air sampler has been benchmarked as an efficient high flowrate sampler for bioaerosol studies.^17,18^ We determined the minimum detectable cell number using our cell recovery and DNA extraction procedure from SASS 3100 standard filters using an *IMP-4* positive *Enterobacter hormaechei* isolate. Spiking at defined concentrations, this pathogen was consistently detectable across all replicates at a quantity of 5000 cells per filter (**Figure 1A, 1B**). This was the limit of detection by both quantitative PCR (targeting the 16S rRNA V4 gene region) and standard PCR (targeting the *IMP-4* carbapenemase gene). Lower quantities were occasionally detectable in some replicates, indicating it is possible but not reliable to observe more dilute signals. Under our standard sampling conditions of 300 L min^-1^ for 4 hours (a total of 72 m^3^ of air per filter), we therefore expect to quantify and reliably distinguish genuine airborne microbial signals from contaminants with as few as 69 target cells per m^3^ of air.

**Figure 1:**
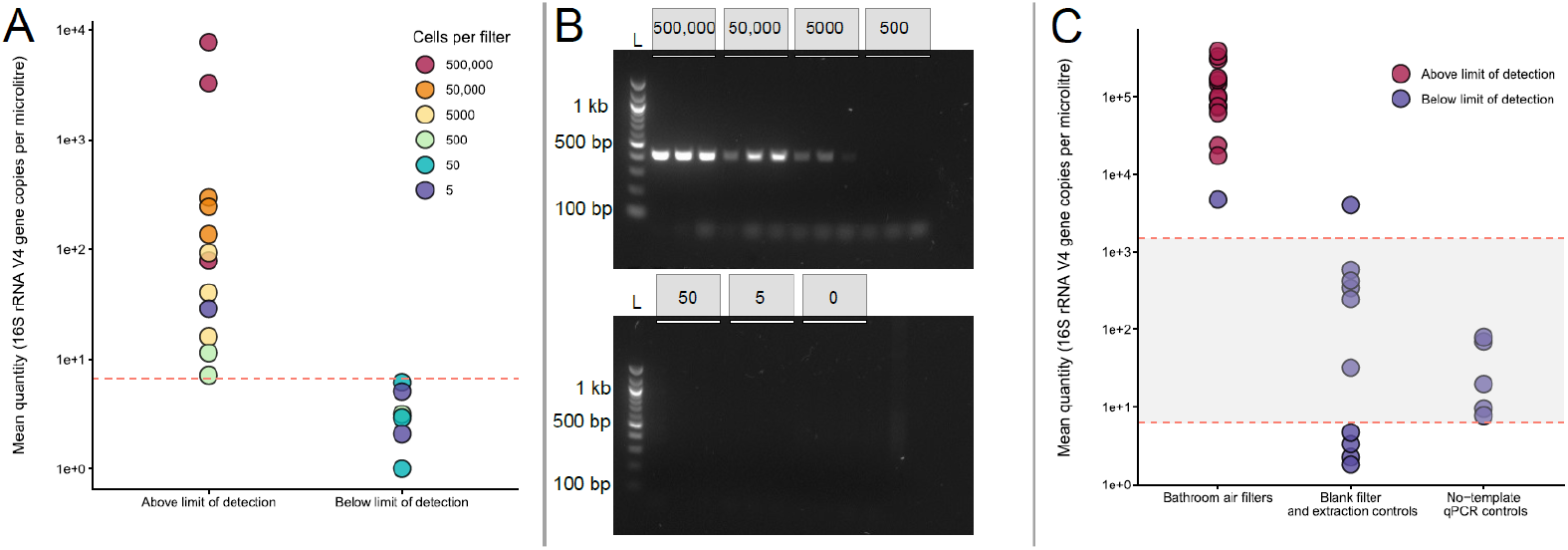
Limit of detection on air filters by quantification of DNA. Defined amounts of *E. hormaechei* cells were spiked onto SASS 3100 standard filters. Triplicate samples spiked with 5000 cells per filter were consistently detectable above blanks by A) 16S rRNA V4 qPCR, where the red dotted line represents the no-template control (NTC), designated as the limit of detection (approx. Cq = 35), and B) standard PCR targeting the carbapenemase gene *IMP-4*. C) Applying a uniform detection threshold of Cq < 35, air samples collected in a workplace bathroom were consistently quantifiable and distinguishable from blank filters, negative extraction controls, and NTCs by 16S rRNA V4 qPCR. The shaded area between the dotted lines represents the range of the Cq = 35 threshold across all qPCR runs when converted to copies per microlitre. Samples are coloured “above” or “below” the threshold for their individual plate, however after correcting for sample dilution, some samples appear above the plotted threshold range.

Based on this limit and the behaviour of no-template controls (NTCs), we designated a Cq value of 35 as an appropriate threshold for detection of microbial biomass by qPCR. Applying this to air samples collected in a workplace bathroom setting, we show that our air sampling and processing approach yields quantifiable biomass that is easily distinguishable from blank filter, extraction, and qPCR controls (**Figure 1C**), increasing confidence that sufficient microbial biomass has been captured to minimise the influence of contaminating DNA in downstream microbiome analyses. To maximise this recoverable biomass, we examined the effect of combining triplicate filters for DNA extraction. While DNA yields increased with filter pooling (mean 0.257 ± 0.15 ng/µl for n = 3 single filters, mean 0.459 ± 0.18 ng/µl for n = 3 filter pools) it did not increase linearly, indicating diminishing returns for the additional cost and labour associated with conducting and processing replicate samples from multiple SASS 3100 samplers.

As a proxy for a low-level airborne pathogen load against a background airborne microbial community, cells were spiked near the limit of detection onto air filter samples collected in the workplace bathroom environment under the standard sampling conditions (4 hours at 300 L min^-1^). Spiked *E. hormaechei* cells were recognisable by amplicon sequencing, despite limited ability for the technique to classify bacteria beyond genus level (**Table 1, Supplementary File 1**). *E. hormaechei* was only classified to family level (*Enterobacteriaceae*), yet at 4000 cells per filter it was identifiable in the 16S rRNA amplicon data as a minor member of the community (2.53%). Almost absent from the background bathroom aerobiome, this family was detectable in the amplicon data at decreasing relative abundance, corresponding to the serial dilution of spiked *E. hormaechei*. This persisted even at 1000 cells per filter (0.33%), indicating it may be possible to recognise key pathogenic organisms distinct from a more typical microbial community in air samples at a density as low as 14 cells/m^3^ (1000 cells in 72 m^3^ of air), particularly if the pathogen is unique at family or genus level and is consistently absent from baseline samples.

**Table 1:** Airborne pathogens may be recognisable by amplicon sequencing. Relative abundance of *Enterobacteriaceae* (putative *E. hormaechei*) spiked onto collected bathroom bioaerosol filter samples by 16S rRNA amplicon sequencing. The family was absent from negative controls.

| Spiked *E. hormaechei* cells | Relative abundance of *Enterobacteriaceae* (% reads) |
| --- | --- |
| 4000 | 2.53 |
| 3000 | 1.15 |
| 2000 | 1.55 |
| 1000 | 0.33 |
| 0 (bathroom aerobiome only) | 0.003 |

### Indoor airborne microbes come from a variety of sources and are regulated by human activity

The microbiome of air samples collected in a workplace bathroom indicated that indoor air contains a diverse mixture of bacteria derived from skin, gut, vaginal, oral and environmental sources, including some potential opportunistic pathogen genera. Culturable isolates from air filter samples were primarily environmentally ubiquitous species (e.g. *Bacillus spp*., *Pseudomonas spp*., *Micrococcus spp*.), but also included likely skin-associated *Staphylococcus* and *Corynebacterium* species and the gut-associated *Enterococcus spp*. Toilet flush aerosols also yielded a mixture of environmental, skin, and gut isolates, and only one pathogenic species (*Acinetobacter baumannii*). *Staphylococcus argenteus*, a member of the *S. aureus* clonal complex with predicted pathogenic potential,^19^ was isolated from the mirror surface and toilet lid. No other putative pathogens were isolated from the workplace bathroom (**Supplementary File 2**).

Following profiling with 16S rRNA amplicon sequencing, the airborne microbiome was consistent across samples collected on different days, suggesting stability of source contributions to the air, and was distinct from blank controls which contained few taxa and appeared to be stochastic rather than representing a common source of contaminating DNA (**Figure 2**). Following filtering of mitochondrial (likely derived from airborne human cells), singleton and contaminant sequences identified by *decontam*,^20^ major bacterial families in bathroom air included *Moraxellaceae* (predominantly *Enhydrobacter*), *Micrococcaceae* (predominantly *Micrococcus*) and *Corynebacteriaceae* (predominantly *Corynebacterium*). Unfiltered negative controls were dominated by the likely environmental contaminants *Bacillus, Novosphingobium, Tepidimonas* and *Pyrinomonas*, which were absent or negligible in the bathroom samples. Thus, even ultra-low biomass samples can provide reliable and interpretable data as long as they are distinguishable from relevant blank controls.

**Figure 2:**
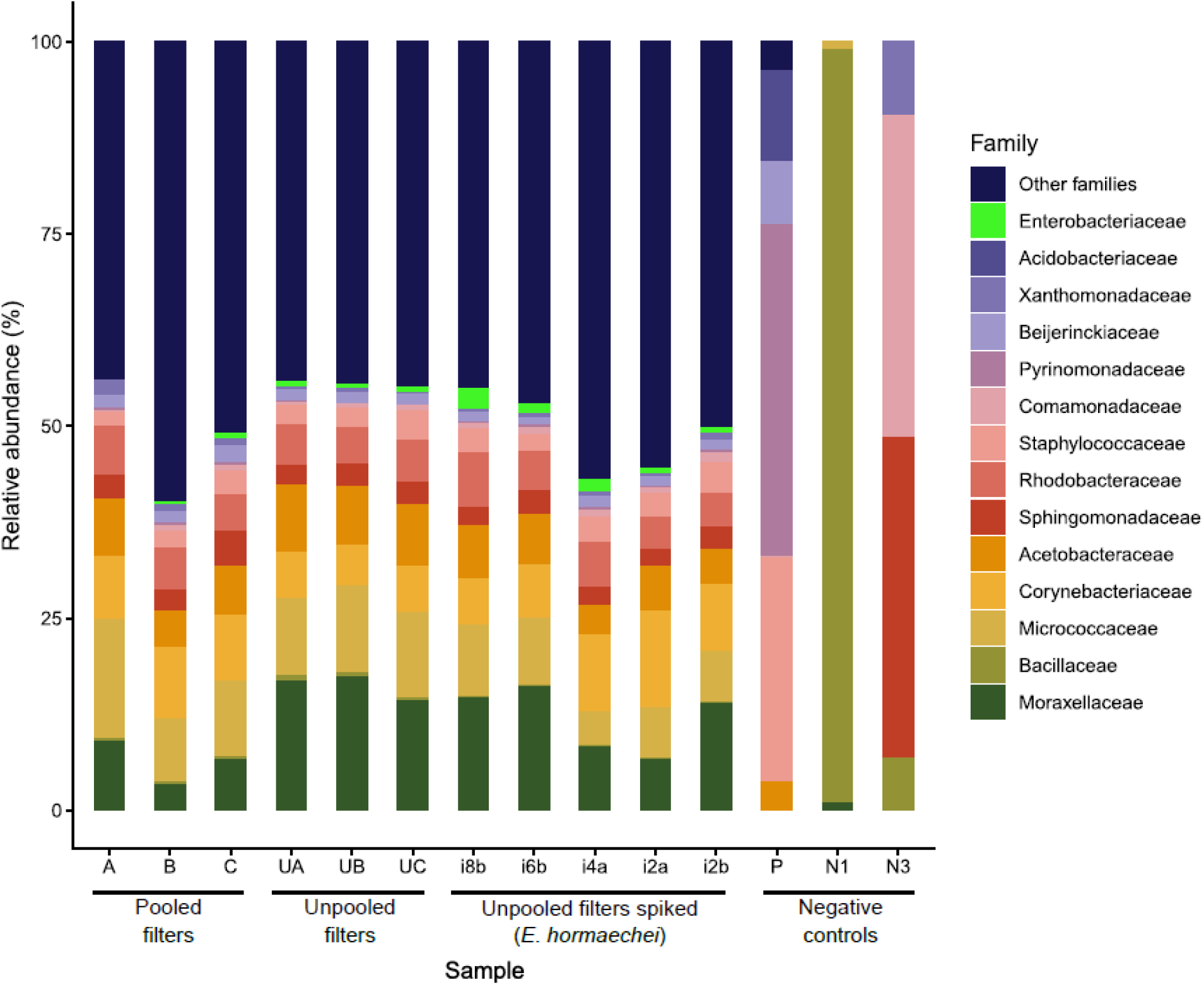
Taxonomic composition of workplace bathroom samples and negative controls. Microbiome profiles are summarised at family level and display the most abundant taxa across all samples (>7% in at least one sample). Samples represent pooled filters (each comprising three filters pooled into one extract), unpooled filters, unpooled filters spiked with *Enterobacter hormaechei* (appearing within *Enterobacteriaceae* shown in bright green), and negative controls (P = passive settling control, N1/N3 = negative DNA extraction controls).

Human activity and occupancy were observed to contribute substantially to airborne biomass (**Figure 3)**. During peak working hours, an average of 4360 cells/m^3^ was recovered from bathroom air samples, whereas this diminished more than two orders of magnitude and below the limit of detection outside working hours to an estimated 9 cells/m^3^ (unpaired t-test, p = 0.0229, **Fig. 3A**). Similarly, the number of detectable cells in the mezzanine lunch area increased when one individual was present (551 cells/m^3^) compared to when the area was vacant (72 cells/m^3^) but the difference was not significant. Supporting this, the diversity of the airborne microbiome (measured by the number of amplicon sequence variants [ASVs] detected) increased with increasing activity in the bathroom (**Fig. 3B**).

**Figure 3:**
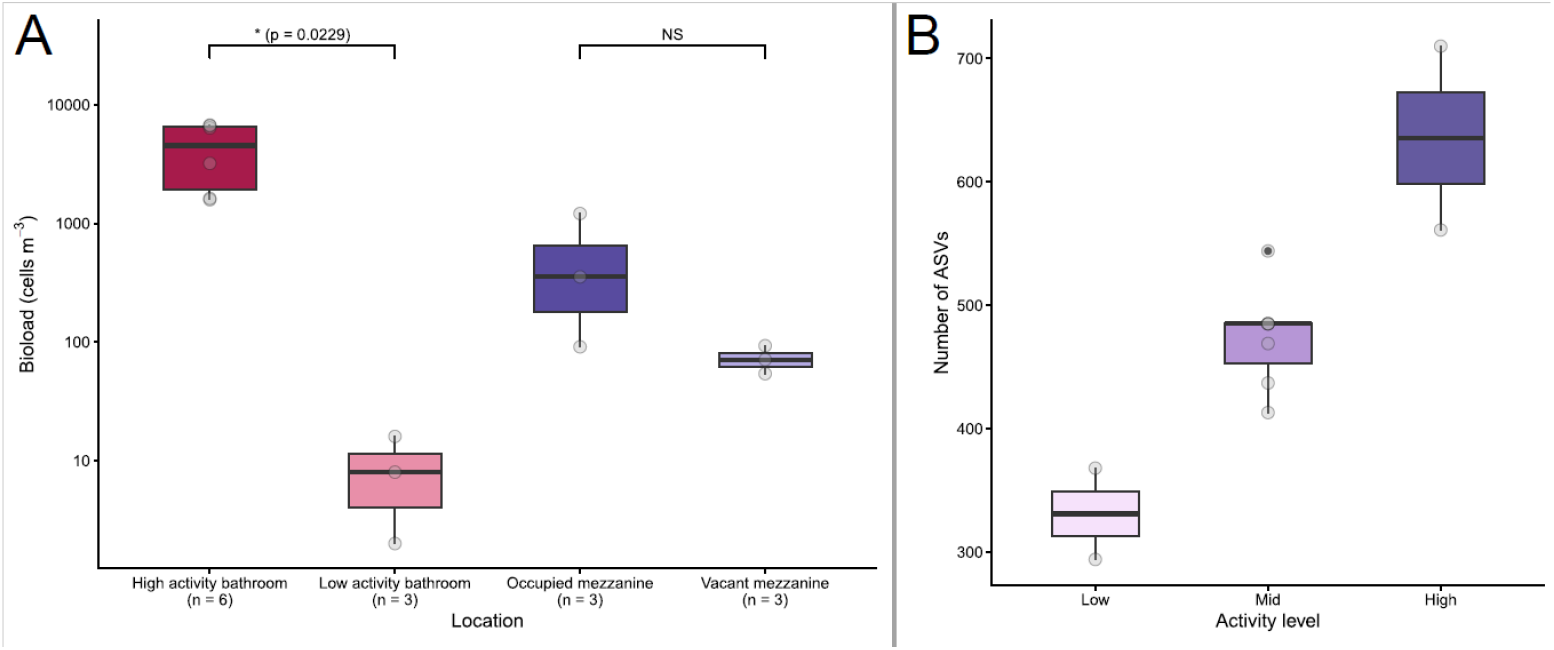
Effect of occupancy and activity on airborne microbial content. A) The airborne bioload in cells per cubic metre, calculated from measured qPCR quantities. B) Community richness measured by number of ASVs from 16S rRNA amplicon sequencing of bathroom filters collected during low (> 1 SD below mean), mid (within 1 SD of mean) and high (> 1 SD above mean) periods of bathroom use, determined by a voluntary activity tally for bathroom use.

### Bacterial strains persist or are re-seeded across bathroom areas

To assess transmission within the bathroom environment, we conducted parallel sampling of toilet flush, toilet lid, mirror and sink surfaces across three separate months (April, June, July) followed by culture-based isolation and identification of bacteria. Isolates derived from the wider bathroom environment were concordant with those found in the airborne community, with seven species isolated from more than one sampling site (**Figure 4A**). *Micrococcus luteus* and three *Staphylococcus* species (*S. haemolyticus, S. warneri* and *S. epidermidis*) isolated from air filters were also culturable from the toilets, indicating a possible source, though *M. luteus* consisted of distinct strains across each area (**Figure 4B**). *S. warneri* was additionally isolated from mirror and sink surfaces, suggesting connectivity between toilet aerosols, the airborne microbiota, and other surfaces. Samples from the mirror yielded few colonies compared to the other sample types, suggesting that dry non-porous vertical surfaces are not a key source of persistent microbes.

**Figure 4:**
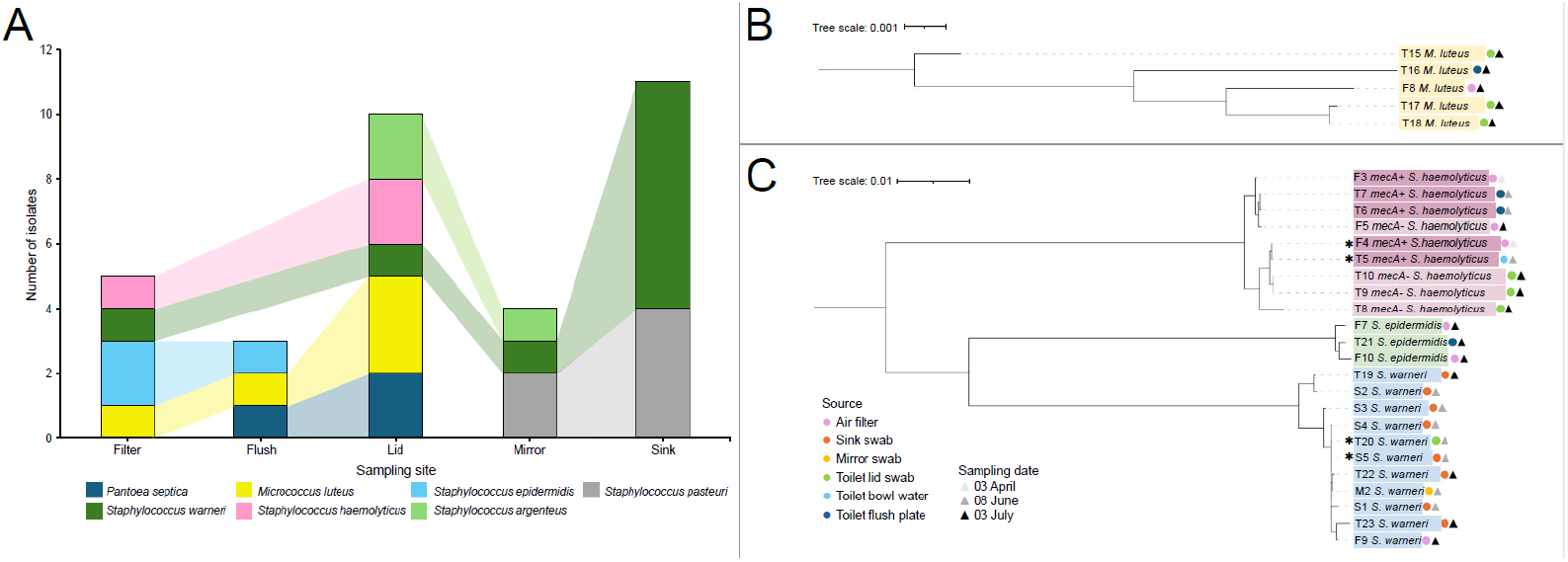
Connectivity between bathroom sources. A) Isolates from the same species were found across sampled bathroom areas. B) *Micrococcus luteus* isolates from different sources were distinct. C) Identical *Staphyloccocus* strains were isolated multiple times over the three-month sampling period. Trees were generated with iqtree and iTOL. Branch lengths are to scale and measured in number of substitutions per site. Species are colour-coded with *Staphylococcus haemolyticus* in pink, *S. epidermidis* in green, and *S. warneri* in blue. Isolates marked with * matched to another isolate to the strain level. Sample source is indicated by coloured circles, and the sampling date by shaded triangles.

At the strain level, whole-genome sequencing revealed evidence of months-long persistence of bacteria across multiple surfaces, including methicillin resistant *Staphylococcus*. Two pairs of identical strains were identified: *mecA*+ *S. haemolyticus* and *S. warneri* (**Figure 4C**). The *mecA*+ *S. haemolyticus* isolates were retrieved two months apart from an air filter in April and toilet bowl water in June, suggesting an environmental reservoir and aerosolised transmission, or a common source in the 2-month interval. The identical *S. warneri* strains were recovered on the same day from a toilet lid and sink swab, suggesting potential travel via human skin shedding or other bioaerosols. While these isolates from a workplace bathroom are likely benign, this demonstrates the potential for connectivity between a toilet source, the airborne microbiota, and bathroom surfaces, with the potential for pathogens and strains harbouring antimicrobial resistance to persist or be reseeded via the air.

### A range of ARGs are naturally present in community settings

The presence of antimicrobial resistant organisms in hospitals is a clear issue, signifying the potential for nosocomial infection with resistant pathogens, or potential emergence of new resistant strains through the exchange of mobile resistance genes. It remains challenging to interpret the risks associated with the presence of antimicrobial resistance alone. In the workplace bathroom setting, we observed a range of antimicrobial resistance determinants detectable by both susceptibility testing of cultured isolates, and by a DNA-based ARG qPCR array. Of the CLSI-recommended Tier 1 antimicrobial agents, six *Staphylococcus* isolates were oxacillin-resistant and two also demonstrated tetracycline resistance (**Table 2**). Other instances of resistance included gentamicin and ciprofloxacin in *Staphylococcus* (Tier 4). Intermediate susceptibility was observed for ciprofloxacin in *Enterococcus* (Tier 2) and ceftriaxone in *Acinetobacter baumannii* (Tier 4). Four of five tested oxacillin-resistant *Staphylococcus* isolates were also cefoxitin-resistant, all of which were *S. haemolyticus*. As oxacillin and cefoxitin are surrogate antibiotics for methicillin, the five oxacillin-resistant *S. haemolyticus* isolates were then confirmed to be *mecA* positive via PCR. The Staphylococcal cassette chromosome mec (SCCmec) is a mobile genetic element permitting the horizontal spread of *mecA* (conferring methicillin resistance) amongst staphylococci. Using SCCmecFinder, *mecA*+ *S. haemolyticus* bathroom isolates did not conform to any of the fourteen recognised SCCmec types and instead demonstrated diverse arrangements (**Supplementary File 2**), demonstrating that despite low biomass, culture-based investigation of the aerobiome can assist with the detection and characterisation of resistant isolates.

**Table 2:** Antimicrobial susceptibility testing of select isolates from a workplace bathroom. TCY (tetracycline), CIP (ciprofloxacin), GEN (gentamicin), IPM (imipenem), CRO (ceftriaxone), OXA (oxacillin), VAN (vancomycin), AMP (ampicillin), LNZ (linezolid), DAP (daptomycin). Bold values indicate resistance, with an asterisk to indicate intermediate resistance. *S. argenteus* uses standards for *S. aureus*. A dash indicates that an MIC was not determined.

| Strain | MICs (µg/ml) |  |  |  |  |  |  |  |  |  |
| --- | --- | --- | --- | --- | --- | --- | --- | --- | --- | --- |
|  | TCY | CIP | GEN | IPM | CRO | OXA | VAN | LNZ | AMP | DAP |
| *A. baumannii* |  |  |  |  |  |  |  |  |  |  |
| T1 (toilet flush) | 2 | 0.5 | 1 | 0.5 | **16\*** | - | - | - | - | - |
| *S. epidermidis* |  |  |  |  |  |  |  |  |  |  |
| F1 (air filter) | **>16** | 0.25 | <0.125 | - | - | <0.0625 | 1 | - | - | - |
| F2 (air filter) | 0.5 | 0.5 | **>16** | - | - | 0.25 | 2 | - | - | - |
| T2 (toilet flush) | 4 | 0.25 | <0.125 | - | - | 0.25 | 2 | - | - | - |
| T3 (toilet flush) | 0.25 | 0.5 | 0.25 | - | - | **>8** | 4 | - | - | - |
| T4 (toilet water) | 1 | 0.25 | 0.5 | - | - | 0.25 | 4 | - | - | - |
| *S. haemolyticus* |  |  |  |  |  |  |  |  |  |  |
| F3 (air filter) | 1 | **>8** | <0.125 | - | - | **>8** | 4 | - | - | - |
| F4 (air filter) | 0.5 | 1 | 0.25 | - | - | **>8** | 4 | - | - | - |
| F5 (air filter) | 0.5 | 1 | <0.125 | - | - | 0.125 | 2 | - | - | - |
| T5 (toilet water) | 0.5 | 0.5 | <0.125 | - | - | **>8** | 1 | - | - | - |
| T6 (toilet flush) | 0.5 | 0.5 | 0.25 | - | - | **>8** | 2 | - | - | - |
| T7 (toilet flush) | **>16** | **>8** | <0.125 | - | - | **>8** | 2 | - | - | - |
| T8 (toilet lid) | 1 | 0.5 | <0.125 | - | - | 0.5 | 2 | - | - | - |
| T9 (toilet lid) | 0.5 | 1 | <0.125 | - | - | 0.25 | 2 | - | - | - |
| T10 (toilet lid) | 0.5 | 1 | <0.125 | - | - | 0.25 | 2 | - | - | - |
| *S. argenteus* |  |  |  |  |  |  |  |  |  |  |
| T11 (toilet lid) | 1 | 0.5 | 1 | - | - | 1 | 2 | - | - | - |
| T12 (toilet lid) | 1 | 0.5 | 2 | - | - | 0.5 | 2 | - | - | - |
| M1 (mirror) | 1 | 1 | 1 | - | - | 0.25 | 2 | - | - | - |
| *E. faecalis* |  |  |  |  |  |  |  |  |  |  |
| F6 (air filter) | - | **2\*** | <500 | - | - | - | 2 | 2 | 0.5 | 0.5 |
| T13 (toilet flush) | - | **2\*** | <500 | - | - | - | 4 | 1 | 1 | <0.25 |
| T14 (toilet flush) | - | **2\*** | <500 | - | - | - | 4 | 2 | 1 | 0.5 |

Assessing community-wide ARG presence with a culture-independent approach, we pooled DNA from nine air filters to input into the QIAGEN microbial DNA array to detect 87 different ARGs. While the sample pool was below the recommended minimum DNA input, we demonstrate that it is possible to confidently detect ARGs by this approach in low biomass air samples. Seven ARGs were detected (OXA-51 group for β-lactamases; *ermB, ermC, mefA, msrA* for macrolide resistance; *tetB* for tetracycline efflux pump; and *mecA*), of which four were detectable at an earlier Ct value than a 5-fold dilution of the array’s positive control DNA. This suggests that air samples amassing a total DNA input comparable to this diluted control, or approximately 40 ng, may be profiled using the QIAGEN ARG array. This indicates that ARGs are identifiable in the airborne microbiome by targeted DNA-based assays even when biomass is low, though extensive pooling is required. These arrays may be practical alternatives to metagenomics, where it remains challenging to collect sufficient airborne biomass to conduct quality sequencing with sufficient depth to detect ARGs, additionally complicated by the presence of human DNA. Establishing a baseline of detectable ARGs may be informative in hospital environments to recognise the emergence of new ARGs or provide an understanding of existing resistance to key antibiotics in use.

### Current air management protocols are effective in a hospital ward housing high-risk patients

We applied our sensitive air sampling approach in patient rooms within a haematology ward, which are subject to a minimum of 6 air changes per hour with HEPA-filtered air inflow. In both occupied and empty patient rooms, minimal biomass was recoverable from the air despite pooling three replicate filters. None of the air filter samples were quantifiable by Qubit, and both fungal (ITS1) and bacterial (16S rRNA, V4) DNA was below our previously defined limit of detection by qPCR for all samples (a Ct value above 35, or above the no-template control for that assay). All hospital air samples were thus indistinguishable from negative controls. Amplicon sequencing of these indistinguishable samples yielded minimal data, but this was higher for the occupied rooms (mean 18,048 denoised and quality-filtered reads per sample) than the unoccupied room (4209 reads) and blank filter control (1459 reads). This corresponded with higher diversity in the occupied rooms, yet the most abundant taxa included sequences affiliated with chloroplasts, mitochondria, and *Deinococcus*, a known laboratory contaminant (**Figure 5**). Following removal of chloroplasts, mitochondria, and sequences identified as likely contaminants due to their presence in the filter extraction control, remaining taxa were affiliated with human skin. *Staphylococcus, Corynebacterium* and *Micrococcus* were present in the occupied patient rooms at an average of 18.17% combined relative abundance (**Table 5**), and only at trace amounts of 0.81% in the empty hospital room. As common skin commensals, *Staphylococcus* and *Micrococcus* are frequently isolated organisms from hospital ward and ICU air,^21,22^ likely introduced into the air by human activity. We do not recommend extensive interpretation of microbiome profiles containing taxa at lower relative abundance than identifiable environmental contaminants and expect such analysis to be misleading in samples that are already below established detection limits.

**Figure 5:**
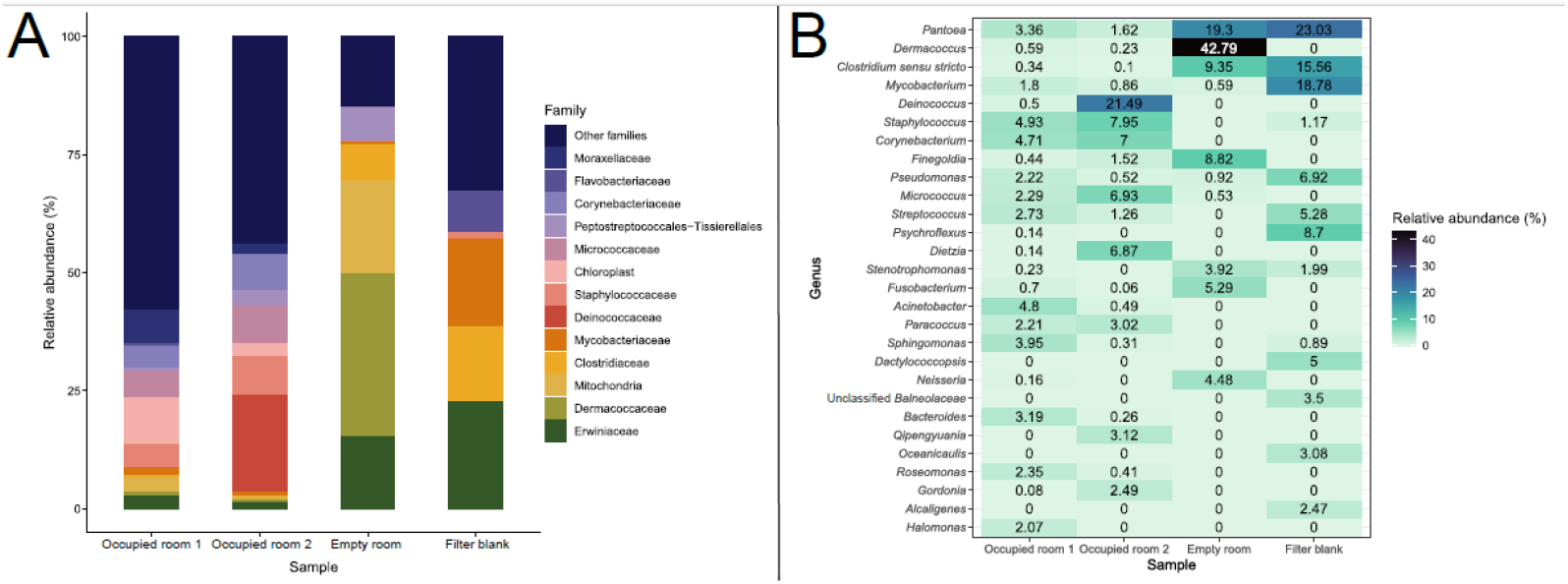
Composition of the hospital aerobiome. Composition of the A) major families, prior to the removal of contaminant mitochondria and chloroplast sequences, and B) major bacterial genera across the samples and controls.

While we cannot declare that the air in these hospital rooms is sterile, combining the quantitative and amplicon sequencing data indicates that no microbes could be reliably detected in hospital air, supporting the efficacy of current measures for air filtration and exchange. The definition of thresholds for confident microbial detection is important when considering the use of airborne surveillance to monitor emerging outbreaks of disease. While amplicon data illustrates the potential of an airborne community in occupied rooms, findings from the empty patient room indicate that this does not constitute an airborne signature that persists.

## Discussion

Despite stringent infection prevention measures, there is an ongoing risk of airborne transmission of infectious diseases in indoor settings such as hospitals, aged care facilities, schools, passenger travel, and workplaces, that is not routinely assessed. Here we developed a method for sensitive detection of airborne pathogens in indoor air environments and tested the approach in a workplace bathroom setting. We found that airborne biomass is strongly associated with human activity, and the airborne microbial community contains a variety of skin, environmental, and gut-associated bacteria, some of which are aerosolised by toilet use and may persist on bathroom surfaces. Testing our approach in individual patient rooms of a haematology ward, we revealed that existing hospital control measures (HEPA filtration and a minimum of six air changes per hour) are effective, as we recovered negligible airborne bacterial signal, indicating that airborne bacteria in these rooms were below our established limits of detection. Amplicon sequencing revealed a small contribution from skin-associated organisms in occupied rooms that was absent from the empty patient room, which are routinely disinfected before housing the next patient. However, these signals were comparable to known reagent contaminants and environmentally ubiquitous organisms such as *Deinococcus, Micrococcus* and *Dietzia*. Our data suggests minimal risk for the transmission or persistence of bacterial pathogens and antimicrobial resistance via the air in individual patient rooms when ventilation is adequate: however, the transmission of viral pathogens, and the aerobiome of communal rooms or facilities are likely to pose an increased risk.

There has been increasing precedence for concern over the risk of bacterial and ARG transmission via indoor hospital air. Outbreaks of MRSA have been recorded in instances where ventilation systems have failed or malfunctioned^23,24^ and recent microbiome-based assessment of hospital bioaerosols has revealed the prevalence of a range of ARGs, ^12,25^ in some cases correlating with resistant infections in the same ward.^12,21,26^ Indoor air microbiome studies have strongly established that human presence and activity is a major driver of bioaerosol density,^27,28^ as we observed in our assessment of the airborne microbial community in a workplace bathroom. However, it has been suggested that adequate ventilation in hospital wards attenuates this, where higher air exchange rates reduce the association between occupancy and culturable bacterial cells from the air.^29^ Hospital bioaerosols may arise from multiple sources, including toilet flushing,^30^ sink drain biofilms that may additionally harbour ARGs,^31^ medical procedures and cleaning,^21^ patient washing,^32^ respiratory expulsions, vomiting, and diarrhoea, or even disturbance of curtains.^33^ Despite these inputs, aerosol residence times are typically short, often under an hour when assisted by ventilation,^34–36^ also reflected in our observations of minimal biomass in the workplace bathroom outside working hours. In our assessment of the hospital air microbiome, while the biomass in air samples was negligible, we observed subtle traces of human activity, with 228 species-level taxa in occupied rooms compared to 13 in the empty patient room. Notably, activities expected to increase aerosolisation, such as cleaning in the empty room and patient toilet use in one of the two occupied rooms, did not produce a measurable increase in airborne bacterial load or enrichment of gut-associated taxa.

Importantly, the extremely low biomass recovered from hospital air highlights a key limitation that similar studies may face, which is that with adequate ventilation, retrievable bacterial signals may be below the limit of detection and are thus highly susceptible to contamination. For example, the dominance of *Deinococcus antarcticus*, a likely contaminant, was present at 20.75% relative abundance in one occupied room, suggesting that background contamination is sufficient to overwhelm genuine signals. The challenges of studying the microbiome of low biomass environments, particularly air, remain present^37^ but are being overcome with improving standards,^14^ such as extensive decontamination and PPE measures for sampling, and careful inspection of blank controls as implemented here. Establishing limits of detection in these scenarios is important to avoid over-interpretation of microbiome data, particularly when *in silico* decontamination methods such as decontam^20^ rely on, and perform best with, samples that are distinguishable from blank controls. Our findings from the workplace bathroom demonstrate that recovery of bacterial biomass from the air is achievable, but reinforces the importance of establishing a limit of detection to appropriately interpret airborne microbiome data, particularly when samples are at or below this limit. Together, this reinforces that airborne bacterial load in individual patient rooms is extremely low under standard infection control methods, which include mandatory N95 masks for medical staff and visitors, minimum ventilation rates of six air changes per hour, air filtration, and pressure differentials.

Based on our findings, it is expected that hospital air samples collected in locations with sufficient ventilation and air filtration measures will commonly contain levels of bacteria below quantifiable limits of detection, but yield DNA sequencing data. Studies that have previously characterised the microbiome or metagenome of low biomass hospital air likely sequence a mixture of background contaminants and genuine biological signals, as has been observed in similar applications like the placental microbiome.^38,39^ However, it is also plausible that airborne transmission of pathogens and AMR occurs transiently and below the limit of detection, yet at sufficient quantities to pose a risk to patients, which is difficult to observe with microbiome approaches. As broad or untargeted detection is key to identifying outbreaks before they occur, it is therefore important to combine these approaches with complementary measures of surveillance to properly identify actionable risks. We showed that the broad qPCR ARG array approach can be effective with low biomass samples that are below the recommended input amount, but importantly these samples needed to be extensively pooled to maximise this input. Broad array approaches can assist with the identification of ARGs new to a given environment, alongside establishing the ‘baseline’ diversity of ARGs that may be present in human commensals or ubiquitous environmental organisms and pose little risk, and guide targeted surveillance efforts. Culture-based surveillance remains valuable for confirming viability, antimicrobial resistance phenotypes, and connecting AMR to hosts.

## Conclusion

In this study, we demonstrate that the indoor aerobiome can be sensitively characterised using optimised contamination-aware sampling approaches, enabling detection of both low-abundance airborne bacteria and antimicrobial resistance determinants. In community settings, airborne microbial load was strongly influenced by human activity and demonstrated connectivity between bathroom areas. These findings highlight that indoor bioaerosols are dynamic and driven by human occupancy. Applying our approach in a hospital ward revealed extremely low airborne biomass, indistinguishable from background contamination, demonstrating the efficacy of stringent air replacement and filtration measures in minimising bioaerosols in patient rooms. However, the absence of a measurable microbiome does not equate to the absence of risk, with regular but transient release of bioaerosols in hospitals from toilets, sinks, and people. We recommend complementary approaches including culture-based methods to detect potential transmission threats and establishing limits of detection and careful interpretation of microbiome data from samples at or below these limits. Overall, maintaining effective air control remains essential and effective for reducing airborne transmission risk, particularly in high-traffic areas or rooms with known sources of aerosol generation. Incorporating periodic airborne surveillance in these areas is likely to provide valuable insight into potential transmission pathways and support outbreak investigations.

## Materials and methods

### Bioaerosol sampling and limit of detection assays

Air samples were collected with SASS 3100 Dry Air Samplers (Research International). Three units were run in tandem operating at their maximum flow rate of 300 L/min for 4 hours, collecting a total of 72 m^3^ of air per filter (standard electret filters). To minimise sample contamination, we implemented decontamination procedures and the use of personal protective equipment (PPE). The SASS 3100 samplers were disassembled prior to each sampling location. Covers and screws were soaked in 5% bleach for 5-10 mins, rinsed with Milli-Q water, and allowed to air dry. Components were then wiped with 70% ethanol and DNA Away Surface Decontaminant (Thermo Scientific) before re-assembly. Gloves, face mask, a full-body disposable coverall and safety goggles were worn during sampling setup and retrieval to minimise contamination from the operator. Once placed, the air samplers were decontaminated with 70% ethanol followed by DNA Away and fitted with a sterile filter. Active air intake commenced following a programmed delay to allow the operator to exit the room.

Air sampling was conducted at two sites: the Monash Biomedicine Discovery Institute (a workplace bathroom setting, and an open mezzanine area) and The Alfred Hospital (a hospital ward setting). In the workplace setting, permission was obtained from facilities managers, and staff and students were notified of the project’s intent and provided with investigator contact details. In the hospital, ethics approval was obtained from The Alfred Hospital Ethics Committee (Project No. 368/24). The bathroom site contained five toilet cubicles, three handwashing sinks, an electric hand-dryer and a paper towel dispenser. Filter blanks involved SASS filter setup and exposure to air outside the bathroom, followed by immediate collection, with the sampler remaining off. For passive air settling controls, the filter was left in place for 5 minutes before removal. In The Alfred Hospital, sampling occurred in single patient rooms in the 7 East haematology ward. These patient rooms were subject to a minimum of six air changes per hour with HEPA-filtered air inflow. All nurses, medical staff, cleaners, and visitors were required to wear a fitted mask whilst in the ward. Samples were taken in an empty patient room and two different occupied patient rooms, with samplers mounted on tripods and the inlet facing the patient bed.

Filters for bacterial culture were stored at ambient temperature for elution and plating on the same day. Filters for DNA extraction were stored frozen at −20ºC within 1 hour of collection prior to DNA extraction.

To evaluate the limit of detection of the SASS 3100 sampler and filters with our modified CTAB DNA extraction protocol, a clinically relevant nosocomial pathogen was serially diluted and inoculated on SASS filters. *Enterobacter hormachei* strain CPO092^40^ contains the IMP-4 gene encoding carbapenemase and was used as a representative hospital-acquired infection agent. An overnight culture of the strain in LB broth was spiked onto blank sterile SASS 3100 filters by pipetting two drops of 2.5 μL culture diluted in PBS, to inoculate a total of 5, 50, 500, 5000, 50,000, and 500,000 cells per filter with three technical replicates. Filters were dried inside a biosafety cabinet until no drops were visible. CFU plate counts were used to confirm accurate cell dilution and delivery. Previously collected bathroom air sampling filters were also spiked with 4000, 3000, 2000, and 1000 cells per filter in duplicate to examine sensitivity with a background microbial community.

### DNA extraction from SASS 3100 filters

Air filters were extracted using a CTAB DNA extraction protocol optimised for low biomass air samples, adapted from Warren-Rhodes *et al* (2019).^41^ Extractions were performed in biosafety cabinets cleaned with 70% ethanol and DNA Away and UV sterilised. Equipment and materials were wiped with ethanol and DNA Away and subjected to UV sterilisation. Reagents were prepared fresh from dedicated stocks and autoclaved or filter-sterilised where appropriate to minimise contamination. Negative controls consisted of both filter and no-filter extraction controls, whereby a sterile filter or just the reagents were processed in the same manner as a filter sample.

Filters were sliced into quadrants with single-use scalpel blades in sterilised petri dishes and transferred into 15 mL TeenPrep Lysing Matrix J bead beating tubes (MP Biomedicals), with forceps sterilised with ethanol and DNA Away samples. Per 15 mL bead tube, 1920 μL of 100 mM phosphate buffer and 1920 μL of SDS lysis buffer was added, vortexed, homogenised in the FastPrep-24 homogeniser (MP Biomedicals) at 5.5 m/s for 30 s, and centrifuged at 4816 x g for 5 min. CTAB buffer (1380 μL) with freshly-added 0.4% β-mercaptoethanol was added to the tube and vortexed prior to incubation 60°C with shaking for 30 min. Tubes were centrifuged at 4816 x g for 1 min at 4°C. In a fumehood, 3300 μL chloroform:isoamyl alcohol (24:1) was added. Tubes were then vortexed for 15 s, chilled for 3 mins at 4°C, and centrifuged at 4816 x g for 10 min at 4°C. The upper aqueous phase was transferred to a 15 mL DNA lo-bind tube, where an additional 3300 μL aliquot of chloroform:isoamyl alcohol was added and the phase separation repeated in samples where the aqueous and inorganic phases were not distinguishable. In a biosafety cabinet, 10 M ammonium acetate was added to the retrieved aqueous phase to a final concentration of 2.5 M, vortexed, and centrifuged at 4816 x g for 5 min at 4°C. The upper aqueous layer was transferred to a new 15 mL DNA lo-bind tube, 0.54 volumes of isopropanol added, mixed by inversion, and incubated at −;20°C for 24 h. Samples were centrifuged at 16,800 x g at 4°C for 20 min to pellet DNA. The supernatant was carefully removed and the pellet was washed and resuspended in 1 mL 70% ethanol, collected again at 16,800 g at 4°C for 20 min, washed once more without resuspending and finally collected (16,800 g at 4°C for 10 min) and supernatant removed. The pellet was dried with a 60°C heat block in a biosafety cabinet for 10-15 minutes, and resuspended in 20 μL microbial DNA-free water. DNA was quantified with the Qubit dsDNA HS assay kit and the Qubit 2.0 fluorometer (Life Technologies).

### Quantitative PCR

qPCR quantification was performed on a QuantStudio 7 Flex Real-Time PCR machine (Applied Biosystems). Biomass was quantified via qPCR targeting prokaryotes (16S rRNA V4 region with primers 515F 5’-GTGYCAGCMGCCGCGGTAA-3’ and 806R 5’-GGACTACNVGGGTWTCTAAT-3’), and fungi (ITS1 region with primers ITS1-F 5’-CTTGGTCATTTAGAGGAAGTAA-3’ and 5.8S 5’-CGCTGCGTTCTTCATCGA-3’). Each reaction consisted of 9 μL of master mix (Roche LightCycler SYBR Green I Master, 0.4 μM each primer) and 1 μL sample DNA. Standards (the 16S rRNA V4 region inserted in a pMA plasmid, and a gBlock containing the ITS1 region from *Candida albicans*) were serially diluted to the range of 10^8^ through to 10^1^ copies/µL and run in duplicate. DNA extracts were diluted either 1:10 or 1:100 to minimise inhibition from salts remaining from DNA extraction, and were measured in triplicate. PCR grade water was used as a no template control. Cycling conditions for 16S rRNA V4 assays involved pre-incubation at 95°C for 3 min and 50 cycles of 95°C for 30 s, 54°C for 30 s, and 72°C for 24 s. For ITS1, the annealing condition was 53.5°C for 30 s and extension at 72°C for 27 s.

### Polymerase chain reaction for IMP-4 gene

Polymerase chain reaction (PCR) amplifications were performed on a Mastercycler Nexus GX2 machine (Eppendorf). A standard KOD Hot Start DNA polymerase PCR reaction contained 1X buffer, 1.5 mM MgSO_4_, 0.2 mM dNTPs, 0.3 μM of each oligonucleotide, 1 unit of KOD Hot Start DNA polymerase (Merck), ~150 ng template DNA, and PCR grade water to make up to 50 μL. Cycling conditions were 95°C for 2 min 35 cycles of 95°C for 20 s, the lowest Tm°C of the oligonucleotides for 10 s, and 70°C for 10 s per kb of expected product; and a hold stage at 12°C. The IMP-4 gene was amplified with primers AP1632 (5’-GGCGTTGTTCCTAAACATGG-3’) and AP1633 (5’-GTGGGTTTAATGGATCTGGC-3’).^42^

PCR products were visualised on 1.2-2.0% w/v agarose gel made with 1X TAE buffer with 2 μL of SYBR Safe DNA gel stain per 50 mL agarose gel (SeaKem LE agarose, Lonza Bioscience). Reference markers were 100 bp or 1 kb ladders (New England Biolabs). Samples were loaded by mixing 5 μL with 2 μL of 6X purple gel loading dye (New England Biolabs), electrophoresed at 100 V and 400 mA for 30-45 min, and visualised with a ChemiDoc Imaging System (Bio-Rad).

### Bacterial isolation and antimicrobial susceptibility testing

Air filters were sliced into quadrants, transferred into screw capped tubes, vortexed at 2500 rpm for 1 min with 7 mL 0.1% Triton X-100, and sonicated in a Powersonic 410 water bath sonicator at the maximum setting of 3 for 1 min to dislodge cells from the filter matrix. Vortexing and sonication steps were repeated to a total of three cycles. Samples were then centrifuged and plated as lawn cultures on BHI agar with and without 1 μL/mL amphotericin. Plates were incubated at 37°C overnight in either aerobic or anaerobic conditions, or 25°C aerobically. Colonies of diverse morphologies were selected and streaked to single colonies with the aim of isolating different organisms and stored as 25% glycerol stocks.

Toilet flush plates were collected from the female bathroom by attaching two agar plates to the inside of the toilet lid. The toilet was flushed once, and bioaerosols allowed to settle for 30 s before retrieval of plates. Toilet water (50 ml) was collected and centrifuged at 8000 x g for 5 min, and the pellet resuspended and plated onto BHI and R2A agar. BHI agar plates (Difco, BD) were incubated at 37°C for 24 or 48 h until colonies were observable. R2A agar plates were incubated at 30°C for 3 to 5 days until colonies were visible.

MALDI-TOF of filter isolates was performed using the direct transfer method on a MALDI Biotyper (Bruker). Isolates were retained for further analysis if the MALDI ID score was greater than 2.0, or if the ID score was greater than 1.7 and the best and 2nd best matches were consistent.

### Minimum inhibitory concentration (MIC) testing

Antimicrobial susceptibility of select bathroom isolates identified as potential human pathogens was assessed using the CLSI broth microdilution method (M07 12^th^ ed.).^43^ Briefly, this involved 96-well plates with antibiotics serially diluted 1:2 in 100 μL of Cation-Adjusted Mueller-Hinton Broth (BD), or 50 mg/L Ca^2+^ Mueller-Hinton broth for daptomycin, or 2% NaCl-supplemented CAMHB for oxacillin. Each well was inoculated with 10 μL of 5 x 10^6^ CFU/mL suspension and observed at 16-24 h depending on antibiotic and organism. QC strains and antibiotics for testing were selected based on CLSI recommendations and inoculated as previously described, to confirm correct preparation of antibiotic dilutions by referring to the expected MICs for the QC strain and antibiotic. Interpretations of susceptible, intermediate and resistant were made using the CLSI guidelines. (M100 Ed36).^44^

### Disk diffusion assay

Oxacillin-resistant isolates were further tested by cefoxitin assay. Overnight cultures of oxacillin-resistant isolates were resuspended in heart infusion broth to 1 x 10^8^ CFU/mL and swabbed as a lawn onto Mueller Hinton agar (BD). 30 μg cefoxitin disks (Oxoid) were then placed onto the surface. All isolates tested were *Staphylococcus* spp. and observed at 24 h. Isolates resistant to both oxacillin and cefoxitin were selected for DNA extraction and screened for the *mecA* gene by PCR using the universal *mecA* primers (forward 5’-ACGTTACAAGATATGAAG-3’, reverse 5’-ACATTAATAGCCATCATC-3’). Interpretations of susceptible, intermediate and resistant were made using the CLSI guidelines (M100 Ed36).^44^

### Antimicrobial resistance gene qPCR array

The QIAGEN 384-well microbial DNA qPCR array was used to assess the feasibility of larger-scale ARG identification arrays for low biomass air samples. A reaction mix comprised 510 μL microbial qPCR mastermix, 60 μL sample containing ~25 ng DNA, and microbial DNA-free water to a total volume of 1020 μL per sample. The negative control was microbial DNA-free water. The positive control was the QIAGEN-supplied microbial DNA positive control containing oligonucleotide targets for each probe. The positive control was also diluted 1:5 to reflect DNA concentrations obtained from air samples. Cycling conditions were an initial activation of 95°C for 10 min, and 40 cycles of 95°C for 15 s and 60°C for 2 min.

### Nanopore sequencing and mecA typing

mecA+ *S. haemolyticus* and species recovered across multiple sampling sites in the workplace bathroom were extracted with the DNeasy Blood & Tissue Kit (QIAGEN). Nanopore library preparation and sequencing was done according to manufacturer protocol with the ONT Rapid Barcoding Kit 96 V14, run on R10.4.1 flow cells with the Oxford Nanopore MinION Mk1B device, and base-called using MinKNOW software. Raw reads were filtered with Filtlong (https://github.com/rrwick/Filtlong) to retain the top 95% reads based on quality and discarded if below 1 kbp in length, followed by assembly with Flye^45^ using the Clinopore pipeline (https://github.com/HughCottingham/clinopore-nf). Genomes were then taxonomically assigned with GTDB-Tk v2.3.2^46^ using GTDB release 214.^47^ Iqtree v2.3.6^48^ was used to construct a phylogenetic tree with the automatically determined best-fit substitution model LG+F+I+R2, and iTOL^49^ for tree visualisation. SCCmecFinder v1.2^50^ was used to contextualise *mecA* in *mecA*+ isolates, using a 90% ID threshold, 60% minimum length and the default reference database.

### 16S rRNA amplicon sequencing and microbial community analysis

Air filter DNA extracts were processed at the Australian Centre for Ecogenomics (ACE). The 515F (5’-GTGYCAGCMGCCGCGGTAA-3’) and 806R (5’-GGACTACNVGGGTWTCTAAT-3’) primers were used to amplify the 16S rRNA V4 region,^51,52^ and amplicons were sequenced on an Illumina V3 MiSeq 2 x 300 bp run.

Amplicon sequence data was processed with QIIME2 v2024.5.0.^53^ Reads were trimmed to remove primers (qiime cutadapt), and quality filtered, truncated, and denoised (qiime dada2)^54^. Based on where median quality dropped below 30 and with increased trimming for an average sequence retention of 80% where necessary, forward and reverse reads were truncated to 192 and 161 bases for the bathroom dataset, and 200 and 190 bases for the hospital room dataset. Identification and removal of contaminants used qiime decontam^20^ with the frequency-based method and default significance threshold of 0.1. Sequences were then filtered for mitochondrial and chloroplast sequences, singletons, and doubletons (qiime taxa filter-table, qiime feature-table filter-features). Reads were grouped into amplicon sequence variants (qiime feature-table), aligned (qiime alignment), and a rooted phylogenetic tree generated (qiime phylogeny). Bathroom samples were rarefied to 9241 reads to retain experimental samples and exclude controls for alpha and beta diversity analysis (qiime diversity). Classification used a classifier trained with RESCRIPt-processed reference taxonomy^55^ and sequences derived from the SILVA 138 SSURef NR99 database (qiime feature-classifier)^56^.

## Supplementary information

Supplementary File 1: Amplicon sequencing read statistics and ASV tables for the workplace bathroom samples.

Supplementary File 2: List of bacterial isolates obtained from workplace bathroom sampling and identified by MALDI-TOF. MALDI-TOF isolates shown had a MALDI ID score greater than 2.0, or an ID score greater than 1.7 and the best and 2nd best matches were consistent. Typing of the SCCmec element is included for *mecA* positive isolates.

Supplementary File 3: Amplicon sequencing read statistics and ASV tables for the hospital ward air samples.

## Acknowledgements

We acknowledge Matthew Parker, Quynh Doan, Mariel Beiers and Ruzeen Patwa for critical technical assistance.

This work was supported by an Australian Research Council Discovery Early Career Award (DE230100542, to RL) and a Human Frontiers Science Program Grant (RGY0058/2022, to CG). PS is supported by a Biomedicine Discovery Scholarship (Monash Biomedicine Discovery Institute). We are grateful to the Australian Centre for Ecogenomics (University of Queensland) for amplicon sequencing services. This work was supported by Monash eResearch capabilities, including the M3 and MonARCH HPC systems.

